# Grey matter volume changes and corresponding cellular metrics identified in a longitudinal *in vivo* imaging approach

**DOI:** 10.1101/559765

**Authors:** Livia Asan, Claudia Falfan-Melgoza, Wolfgang Weber-Fahr, Carlo Beretta, Thomas Kuner, Johannes Knabbe

## Abstract

Magnetic resonance imaging (MRI) of the brain combined with voxel-based morphometry (VBM) has revealed structural changes of grey and white matter in a range of neurological and psychiatric disorders. However, the cellular basis of volume changes observed with VBM has remained unclear. We devised an approach to systematically correlate changes in grey matter volume (GMV) with cellular composition. Mice were alternately examined with structural MRI and two-photon *in vivo* microscopy at three time points, taking advantage of age-dependent changes in brain structure. We chose to image fluorescently labelled cell nuclei, because these can be readily imaged in large tissue volumes and allow inferences on several structural parameters: (1) the physical volume as determined from a subset of nuclei used to generate a geometrically defined space, (2) the number of cells, (3) the nearest neighbour distance measured between all nuclei as an indicator of cell clustering, and (4) the volume of the cell nuclei. Using this approach, we found that physical volume did not significantly correlate with GMV change, whereas mean nuclear volume was inversely correlated. When focusing on layers within the imaging volume, positive correlations of GMV were found with cell number near the cortical surface and nearest neighbour distance in deeper layers. Thus, the novel approach introduced here provided new insights into the factors underlying grey matter volume changes.

## Introduction

Magnetic resonance imaging (MRI) has tremendously advanced our understanding of brain structure and function in health and disease. Brain morphology and tissue composition can be examined using a variety of structural scanning protocols (Radua, Canales-Rodriguez et al. 2014). A range of automated image analysis tools providing unbiased results have been developed in the past decades to quantify structural changes. One of the most widely used computational approaches is voxel-based morphometry (VBM) (Ashburner and Friston 2000, Salmenpera and Duncan 2005). VBM provides an automated quantitative analysis of the distribution of grey matter (GM) and white matter (WM) to detect voxel-wise differences in brain tissue concentration (e.g. Grey Matter Density, GMD). GMD is then modulated by multiplication with the Jacobian Determinant (JD), resulting in a readout that states grey matter volume (GMV). VBM has been applied to examine physiological aging (Good, Johnsrude et al. 2001) as well as to a vast spectrum of diseases, including depression (Grieve, Korgaonkar et al. 2013, Matsuo, Harada et al. 2019), panic disorder (Uchida, Del-Ben et al. 2008), posttraumatic stress disorder (Kuhn and Gallinat 2013), chronic pain (Apkarian, Sosa et al. 2004), Alzheimer’s disease (Matsuda 2013) and many more. Translational studies have become possible by extending VBM to animal studies, leading to a better understanding of pathology-related changes in brain structure (Seminowicz, Laferriere et al. 2009, Biedermann, Fuss et al. 2012, Bilbao, Falfan-Melgoza et al. 2018). Despite the widespread application of VBM, the physical basis of the terms “tissue concentration” or “volume”, commonly used by VBM studies to describe structural changes, remained poorly defined and mostly serve as semantic wild cards. However, a mechanistic understanding of the physical and cellular basis of GMV changes is pertinent (Figure 1A) (Henderson and Di Pietro 2016, Kuner and Flor 2017, Pomares, Funck et al. 2017).

**Figure 1.**
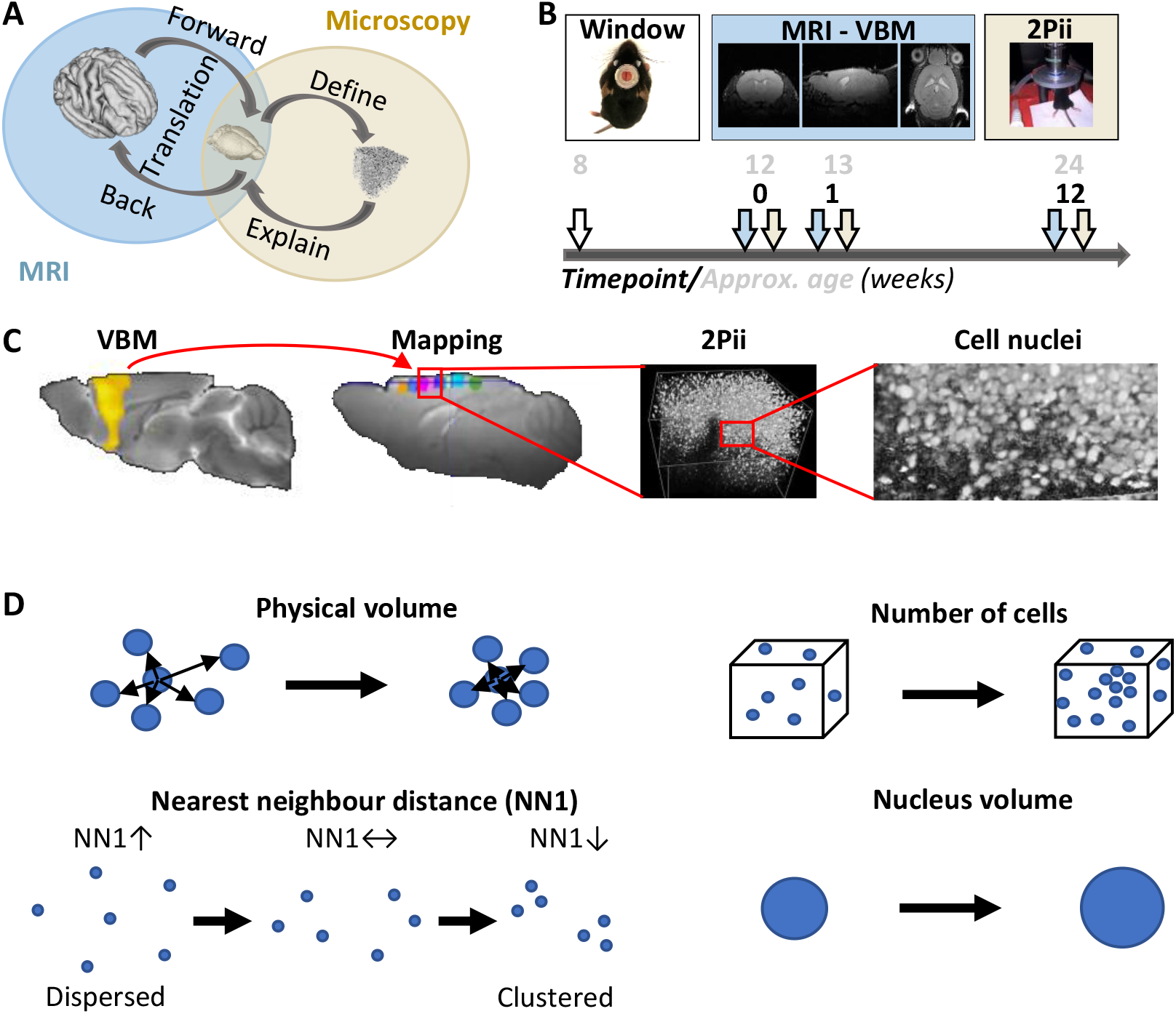
Investigating cerebrocortical structure across species, size and time. **A.** Using MRI in human and rodent studies we can directly translate findings between species with the same imaging modality. Explorative whole brain studies identify key regions which can be further investigated by microscopy studies with mice, revealing specific cellular mechanisms for large-scale structural observations. **B.** MRI and 2Pii allow for repetitive imaging *in vivo* and therefore capture dynamics of brain structure in individual animals. The imaging intervals were precisely defined in each mouse (time points given in black), while the corresponding age of the mice may differ ± 1 week due to the staggered experimental design. Brain images taken from https://scalablebrainatlas.incf.org. **C.** VBM results shown on brain template, mapping of MRI volumes to 2Pii volumes, 3D 2Pii stack and zoomed in view showing the distribution of cell nuclei. **D.** Parameters determined from imaging cell nuclei in 3D space. Further details see main text.

Potential mechanisms underlying VBM volumetric changes may include altered neurogenesis and glial proliferation, changes in neuronal or glial size, angiogenesis and endothelial cell proliferation, shifts in dendritic spine size and density or remodeling of axonal processes and alterations of the extracellular space (e.g. by changes in blood-perfusion and ultrafiltration, edema, glymphatic system) (Lerch, Yiu et al. 2011, Streitburger, Moller et al. 2012, Keifer, Hurt et al. 2015). Furthermore, hormonal changes, medication and a number of other different factors can influence global brain volume on a short-time basis (Dieleman, Koek et al. 2017). To address this lack of mechanistic understanding, studies have been conducted in animals to compare the MRI-detectable volume changes with detailed *ex vivo* histological analyses (Lerch, Yiu et al. 2011, Keifer, Hurt et al. 2015, Biedermann, Fuss et al. 2016). These studies mostly found changes in spine density as well as markers for newborn neurons and astrocytes in the mouse hippocampus.

However, studies comparing *in vivo* VBM results with *ex vivo* cellular assessments typically focus on single time points and lack the ability to follow changes within individual subjects. To overcome these limitations, we designed a longitudinal translational neuroimaging approach (Figure 1B) that combines, in the same set of mice, structural MRI and two-photon *in vivo* imaging (2Pii), a microscopy technique well suited to image superficial cortical volumes through implanted cranial windows (Figure 1B, (Helmchen and Denk 2005)). Corresponding imaging volumes of both modalities were defined by vasculature-based mapping and thereby allowed us to precisely relate changes in VBM to changes in cellular composition (Figure 1C). The latter was obtained from imaging nuclei of all cells in mice harboring nuclei genetically labelled with green fluorescent protein. Several important structural parameters can be inferred from labelled nuclei alone (Figure 1D): (1) the physical volume (i.e. a tissue volume given in µm^3^); (2) their number and therefore the number of all cells in the imaging volume, allowing inferences on cell death, migration or generation of new cells; (3) the nearest neighbour relationship, a composite of physical volume, density of nuclei and spatial cell clustering; and (4) the mean volume of the nuclei. These parameters were individually correlated to the changes in VBM-derived parameters. The known age-dependent changes in brain volume (Good, Johnsrude et al. 2001) was used as a test case to obtain changes in VBM volumes over time and to correlate these with the changes obtained from 2Pii data. Unexpectedly, we did not find changes in the physical tissue volume that could significantly account for GMV changes. Yet, at least part of the GMV changes can be attributed to changes in nearest neighbour distance, amount of cells and volume of cell nuclei.

## Methods

### Ethical approval

This study was carried out in accordance with the European Communities Council Directive (86/609/EEC) to minimize animal pain or discomfort. All experiments were conducted following the German animal welfare guidelines specified in the TierSchG. The local animal care and use committee (Regierungspräsidium Karlsruhe of the state Baden-Wuerttemberg) approved the study under the reference number G294/15.

### Mice

Adult transgenic mice of C57BL/6 background expressing an enhanced green-fluorescent protein fused to human histone H2B (B6.Cg-Tg(HIST1H2BB/EGFP)1Pa/J, short ‘Histone-GFP’, (Hadjantonakis et al. 2004) were purchased from Jackson Laboratories (stock #006069) and bred in our animal facility at IBF (Interfakultaere Biomedizinische Forschungseinrichtung). Since the transgene is controlled by a ubiquitously active CAG-promoter, nucleosomes and chromatin in all cells express the Histone-GFP fusion protein and can be identified by fluorescence imaging. After cranial window surgery, mice were housed separately in individually ventilated cages at a 12-hour light-dark cycle and in a temperature- (22 ± 2 °C) and humidity- (60 % ± 4 %) controlled environment. Food and water were available *ad libitum*. Mice of either sex were used at equal numbers.

### Experimental design

Twelve mice divided into four different batches of three mice each were used. Animals within a batch were littermates. Chronic cranial windows were implanted at eight weeks of age. One animal had to be sacrificed due to window detachment after surgery and dropped out, two others were excluded as bleeding or infection severely compromised image acquisition in 2Pii. Baseline MRI was acquired four weeks after window surgery and followed by 2Pii one or two days later. In the investigated set of mice, we conducted a sham surgery as part of the spared nerve injury model one day after first 2Pii, because these animals will serve as controls in a study related to neuropathic pain. The procedure included a 10 min surgery on the left hind limb under Isoflurane anaesthesia with skin incision, separation of muscles and exposure of the sciatic nerve branches without further intervention before closing the skin. Two consecutive MRI and 2Pii sessions were carried out at one week and at 12 weeks after baseline at consistent times of day. For a visual display of the experimental timeline see Fig. 1B.

### Chronic cranial window surgery

Cranial window surgery was performed as described elsewhere (Knabbe, Nassal et al. 2018). Briefly, mice were accommodated to the surgery room for 30 minutes. 3 µl per gram body weight of a narcotic mix consisting of 120 µl Medetomidine (1 mg/ml, Sedin), 320 µl Midazolam (5 mg/ml, Hameln) and 80 µl Fentanyl (0,05 mg/ml, Janssen) was injected intraperitoneally. Eye ointment was applied, head was shaved, and mouse was placed in a stereotactic head holder when no reaction was shown to hindpaw forceps pinch. Mice were kept on a 32 °C heating pad throughout the procedure. A craniectomy (diameter 6 mm) centered 1 mm cranial to bregma was created with a dental drill. Dura was removed carefully with fine forceps on the exposed part of the right hemisphere to ensure long lasting high image quality. A round glass coverslip of 100 µm thickness and a custom-made light-weight plastic holder ring were positioned on top of the exposed area and cemented to the surrounding skull with dental acrylic, cautiously preventing any glue from touching the brain surface. Coverslips had previously been curved to match natural brain curvature on the mediolateral axis and avoid regional flattening of the brain by adapting them to a 20 mm diameter graphite cylinder by heating them in a laboratory oven (Supplementary Figure 1 (Kim, Zhang et al. 2016)). For mice to recover from anaesthesia after completing the procedure, a mix containing 120 µl Atipamezole (5 mg/ml, Prodivet), 120 µl Flumazenil (0,1 mg/m, Fresenius Kabi) and 720 µl Naloxon (0,4 mg/ml, Inresa) was injected intraperitoneally at 3 µl per gram body weight each. To relieve postsurgical pain, 250 µl of Carprofen (50 mg/ml, Carprieve) was injected subcutaneously immediately after surgery and supplemented every 8-12 h until 24 hours after the surgery. Mice were then placed in home cages on a heating plate and monitored until normal locomotion and grooming behaviour was observed.

### MRI

MR data were acquired in a 9.4 T horizontal bore animal scanner (Bruker, Ettlingen, Germany) with a two-element anatomically shaped cryogenic mouse surface coil cooled to 28 K. The cryogenic coil gives an improvement factor of 2.5–3.5 in signal-to-noise ratio compared to the conventional setup of a 4-channel receiver array and a volume transmit coil. The animals were anaesthetized by a gas mixture of O_2_: 30 % and air: 70 % with 2 % Isoflurane during each measurement. Respiration rate and body temperature (maintained at 36 °C) were monitored throughout the experiment. High-resolution 3D structural images were acquired with a T2-weighted RARE sequence (Rapid Acquisition with Refocused Echoes, RARE factor 16, matrix size 225 x 192 x 96) with an echo time TE = 62.5 ms, a repetition time TR = 1.2 s and a spatial resolution of 78 µm x 78 µm x 156 µm.

### Longitudinal VBM analysis

The acquired structural images were resized by a factor of 10 and co-registered to a template (Biedermann, Fuss et al. 2012) in Paxinos standard space. Pairwise longitudinal non-linear registration was performed for each subject with SPM12 using the one week and 12 weeks - timepoints as comparisons to baseline in order to analyse structural changes. This procedure resulted in three distinct images for each data pair: An average image of the two analysed time points for each subject, a vector field depicting the local shift of each voxel for the registration and an image of the Jacobian determinant describing the local volume change in each voxel between the two time points (Fig. 2A). Afterwards, average images were skull-extracted and segmented, separating grey matter (GM), white matter (WM) and cerebrospinal fluid (CSF) using tissue probability masks respectively (Biedermann, Fuss et al. 2012). The resulting images were multiplied by the Jacobian determinant images in order to generate modulated maps for each tissue class. The DARTEL toolbox (SPM) (Ashburner 2007) was used to create a group-specific tissue class template and normalize all segmented data to a common Paxinos space. The normalized and modulated tissue images depicting the volume difference between two time points were smoothed (4 mm Gaussian kernel) and statistically tested in a second-level (T-test) analysis with SPM12 run under Matlab. While the Jacobian Determinant is the driving mechanism for the detection of longitudinal structural differences within subject changes, segmentation into tissue classes accounts mainly for between-subject difference in local GM distribution. For the analysis of volume changes over time within the 2Pii voxels, therefore values from the Jacobian Determinant images corresponding to each individual 2Pii mask were extracted with the transformed masks and used for correlation analysis.

**Figure 2.**
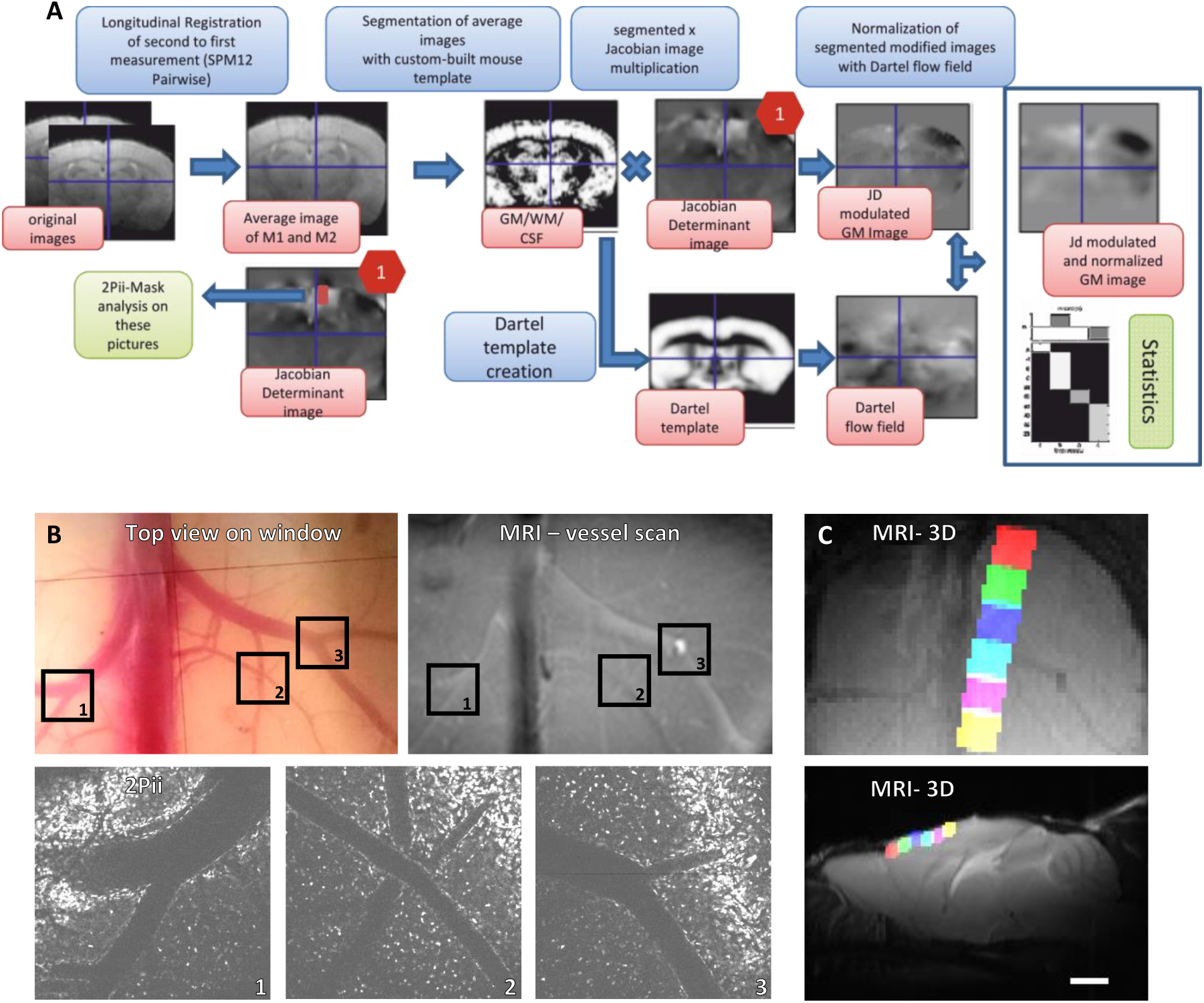
VBM analysis pipeline and cross-modal affine image registration using vessel branchings as fiducial points. **A.** Longitudinal registration processing of MRI 3D images. Longitudinal nonlinear registration was performed for each subject between timepoints, which yielded resulting average and Jacobian Determinant (JD) images. The average images were pre-segmented in three tissue probability maps (GM, WM, CSF) and multiplied by the JD-images. From these a Dartel prior knowledge template was created and the resulting flow fields were used to normalize the JD-modulated tissue maps into paxinos space. They were finally smoothed and statistically analysed in SPM second-level models. **B.** Vessel branchings are visible in MRI as well as 2Pii without any additional contrast agent. **C.** 6 partially overlapping 2Pii-stack-masks of one animal registered on the corresponding MRI. Average intensity projections; up: horizontal view, bottom: sagittal view. Scale bar = 2 mm.

### Cross-modal image registration

To define exact masks that covered the volume of every single 2Pii image stack in the respective MRI space, we exploited superficial venous vessel branching points as natural fiducial landmarks visible in both imaging modalities (Fig. 2B). An individual 2-dimensional affine registration for the XY-positions of the masks was inferred on the MR image for every animal using 3 vessel branching points as fiducials. Z-positions and tilts around all axes were calculated and binary masks fitting the 2Pii-stack dimensions inside the MRI volume were created (Fig. 2C). These binary masks were then transferred to the average image of two compared time points by applying vector fields resulting from longitudinal nonlinear registration (Fig 2D).

### Two-Photon *in vivo* imaging (2Pii)

Mice were acclimatized to the microscopy room for 30 minutes before starting the experiment. Induction of anaesthesia was done with 4-5% Isoflurane (Baxter), then adjusted according to breathing rate at about 0.5-2 % with a O_2_-flow rate of 0.5 l/min. Eye ointment was applied and mouse head was tightly fixated with two metal bars enclosing the head ring to avoid movement artifacts. Two-Photon imaging was conducted with a TriM Scope II microscope (LaVision BioTec GmbH) equipped with a pulsed Ti:Sapphire laser (Chameleon; Coherent) at an excitation wavelength of 960 nm. Emitted fluorescence was passed through a 530/70 nm band pass filter (Chroma) before being collected and amplified a by low-noise high-sensitivity photomultiplier tube (PMT; Hamamatsu, H7422-40-LV 5M). A 16x objective with 0.8 numerical aperture, 3 mm working distance and water immersion (Nikon) was used to scan 3D image stacks with a field of view of 700 µm x 700 µm in XY until a depth of 700 µm below the cortex surface was reached. Image planes were sampled at a step size of 2 µm (voxel size 0.29 µm x 0.29 µm x 2 µm). PMT noise offset was checked and corrected prior to each imaging session to be stable throughout the whole experiment. Increase of laser power was set in defined depths for every mouse individually to assure good signal to noise image acquisition while minimising the amount of oversaturating pixels. Six stack positions were recorded in a straight line immediately lateral to the superficial sagittal sinus from rostral to occipital (with a 15 % inter-stack-overlap), covering parts of the anterior- and midcingulate cortex and secondary motor areas. Stack positions were reidentified at later time points by superficial vessel orientation and aligning image position by nuclei patterns in a depth of ∼200-250 µm below the cortical surface.

### Confocal microscopy of fixed brain slices

Mice were deeply anaesthetized with intraperitoneal Narcoren (500 mg/kg bodyweight) and trans-cardially perfused with 4% paraformaldehyde (PFA) in phosphate-buffered saline (PBS). The brains were post-fixed overnight in 4 % PFA. After washing with PBS, 50 µm thick coronal brain sections were cut with a vibratome. The slices were incubated separately in wells with PBS and 5 ng/ml DAPI (4′,6-diamidino-2-phenylindole) for 20 min before they were rinsed in PBS and mounted in SlowFade Gold (Life Technologies). Superficial cortical areas of anterior and midcingulate cortex were imaged on a Leica SP8 inverted confocal microscope with a 63x oil immersion objective (numerical aperture = 1.4) and a resolution of 0.06μm×0.06μm×0.76μm. Samples were illuminated with a 405 nm laser for DAPI and 488 nm for Histone-GFP in sequential scans. Detectors collected light from 409-464nm for DAPI (PMT) and 491-564nm for Histone-GFP (Leica HyD).

### Microscopic image analysis: Tissue volume changes

To analyse regional cortical volume changes, we identified nuclei patterns in the image stacks that were stable over time, indicating that a nucleus within this pattern did not individually migrate and thus serves as a fiducial marker for a stable position within the stack (Fig. 5A). These marker nuclei were sampled throughout the whole stack. Every change of a volume spanned between markers were consequentially interpreted as an expansion or shrinkage of the tissue volume in between them.

The stack coordinates of the centres of these marker nuclei were collected for every timepoint manually using FIJI (Fig. 5B) (Schindelin, Arganda-Carreras et al. 2012). We used three-dimensional Delaunay triangulation to define tetrahedra whose vertices were given by the point coordinates (Fig. 5C). The volume bounded by the convex hull surrounding all tetrahedra was calculated and was referred to as the ‘physical volume’ in this study. The set of marker nuclei was arbitrarily chosen in each mouse and therefore makes comparisons of absolute volumes difficult. Thus, comparisons were based on a ratio of convex hull volume at one or 12 weeks and the convex hull volume at baseline determined for each mouse separately. A ratio of less than 1 indicates tissue shrinkage, while a ratio larger than 1 indicates expansion during the respective time period. All computations were done with MATLAB (The MathWorks, Inc.).

### Automated image analysis pipeline for nucleus count, nucleus volume and nearest neighbour distances

The bioimage analysis workflow was designed to segment GFP positive nuclei from 2Pii stacks in 3D (Supplementary Figure 2). The 2Pii stacks were deconvolved with Huygens SVI software using the CMLE algorithm, with a SNR of 7 and 500 iterations (Scientific Volume Imaging, The Netherlands, http://svi.nl). To further improve nuclei detection, we used the ilastik autocontext workflow to enhance differentiation from image foreground and background by training the software to recognize varying signal intensities and textures of nuclei as foreground, resulting in a foreground probability map (C. Sommer, C. Strähle et al. 2011). All the acquired z-stacks were processed using the ilastik batch processing option in headless mode operation on the bwHPC cluster with an average running time of 6 h per z-stack.

The 3D nucleus detection and segmentation workflow was implemented as a custom-written MATLAB script that also includes analysis steps in ImageJ/Fiji by using an ImageJ-MATLAB extension (Hiner, Rueden et al. 2017). The workflow involved two major steps: 3D seed detection and 3D watershed segmentation. First, seeds were detected on ilastik probability maps using FIJI’s 2D ‘find maxima’ and ‘3D dilate’ operators and linked as 3D labelled object when connected to each other in z. Secondly, the 3D seeds and the probability maps were used as input for the 3D watershed algorithm with a fixed intensity threshold and assumed nucleus radius. The intensity threshold and the nucleus radius were optimized comparing the output of the automated segmentation with 3D manual segmented ground truth data. To reduce the number of false positive and negative segmented nuclei, a cutoff value for the volume was introduced to the workflow. The cutoff was set according to the nuclei volume between 1500 pixels^3^ and 18000 pixels^3^. The objects below the cutoff value were removed from the analysis while the objects above the cutoff value were stored in a new image stack and re-segmented using a lower 3D watershed threshold value. The new re-segmented nuclei with a volume between the cutoff values were added to the previous segmentation as new labelled objects (Supplementary Figure 2).

The ImageJ-Fiji/Matlab script is fully automated and was used to process all image stacks on a Dell PowerEdge T630 server with 20 cores, 1TB of RAM with an average running time of 3 hours per z-stack. For each input stack, an output folder was created to save 3D-segmented nuclei as labelled objects and a csv file with the xyz nuclei centroid coordinates and the nuclei volume size.

Nuclei centroid data was stored in a SQLite-Database. Z-Densities of all stack centroids were plotted, overlaid for compared timepoints and inspected for large differences. If differences of the two summed up densities of more than 1.0 were present, the corresponding stack image data were manually checked for factors leading to bad image quality, mostly being large areas with missing signal compared to the other timepoints. These could for example be the occasional formation of air bubbles in the immersion water leading to areas without signal within the field of view. Non-comparable image timepoints were excluded from subsequent analysis. In sum, a total of more than 2.5 x 10^6 nuclei were analysed. Ratios of total nucleus count, distances to the nearest neighbour and volume of the segmented nuclei were analysed using the spatstat package in R. (Adrian Baddeley, Ege Rubak et al. 2015).

### Statistics of 2Pii data

Statistical analysis was performed in Prism (GraphPad Software). All distributions were tested for normality and statistical testing was performed accordingly with parametric or non-parametric tests. Details to applied tests can be found in figure subtexts.

## Results

The longitudinal imaging scheme started four weeks after implantation of the cranial windows, when mice were approximately 12 weeks old (see Methods, Fig.1B). A baseline (0 weeks) MRI dataset and a 3D 2Pii dataset were acquired, followed by a second dataset after one week and a third dataset after 12 weeks. Cross modal registration of the volumes was achieved by mapping the blood vessel patterns (see Methods, Fig. 2B), providing a basis to define cubes of 700 µm x 700 µm x 700 µm for quantitative comparison of the two modalities (Fig. 2C).

### Whole brain MRI and VBM analysis

VBM analysis of MRI data revealed areas with increasing or decreasing volumes when comparing timepoints 0 to 1 and 0 to 12 weeks (Fig. 3A, B). During the first week, areas showing volume increases include the cerebellum and midbrain areas while mainly the parietal lobe and parts of the frontal lobe showed volume reduction (Fig. 3, left set of panels). After 12 weeks, the pattern became more pronounced with widespread volume increases in the cerebellum and thalamic areas. In contrast, especially the olfactory bulb, occipital and parietal lobe decreased in volume with most pronounced changes occurring in visual areas (Fig. 3, right set of panels). The quantified volume changes are shown in Figure 3C for 28 atlas regions (Dorr, Lerch et al. 2008). The frontal and prefrontal areas showed strongly lateralized results in this study with a volume loss or lower volume gain compared to the left side. The general pattern of changes, however, was similar to a previous study probing age-dependent GMV changes, with the difference that the mean gain/loss of volume was more shifted to the volume loss which may be due to the different age range investigated in this study (Bilbao, Falfan-Melgoza et al. 2018). In summary, we found age-dependent changes in GMV in distinct areas, including superficial regions of the neocortex that are accessible to 2Pii (Fig. 3A), thereby providing a basis for the correlative approach described here.

**Figure 3.**
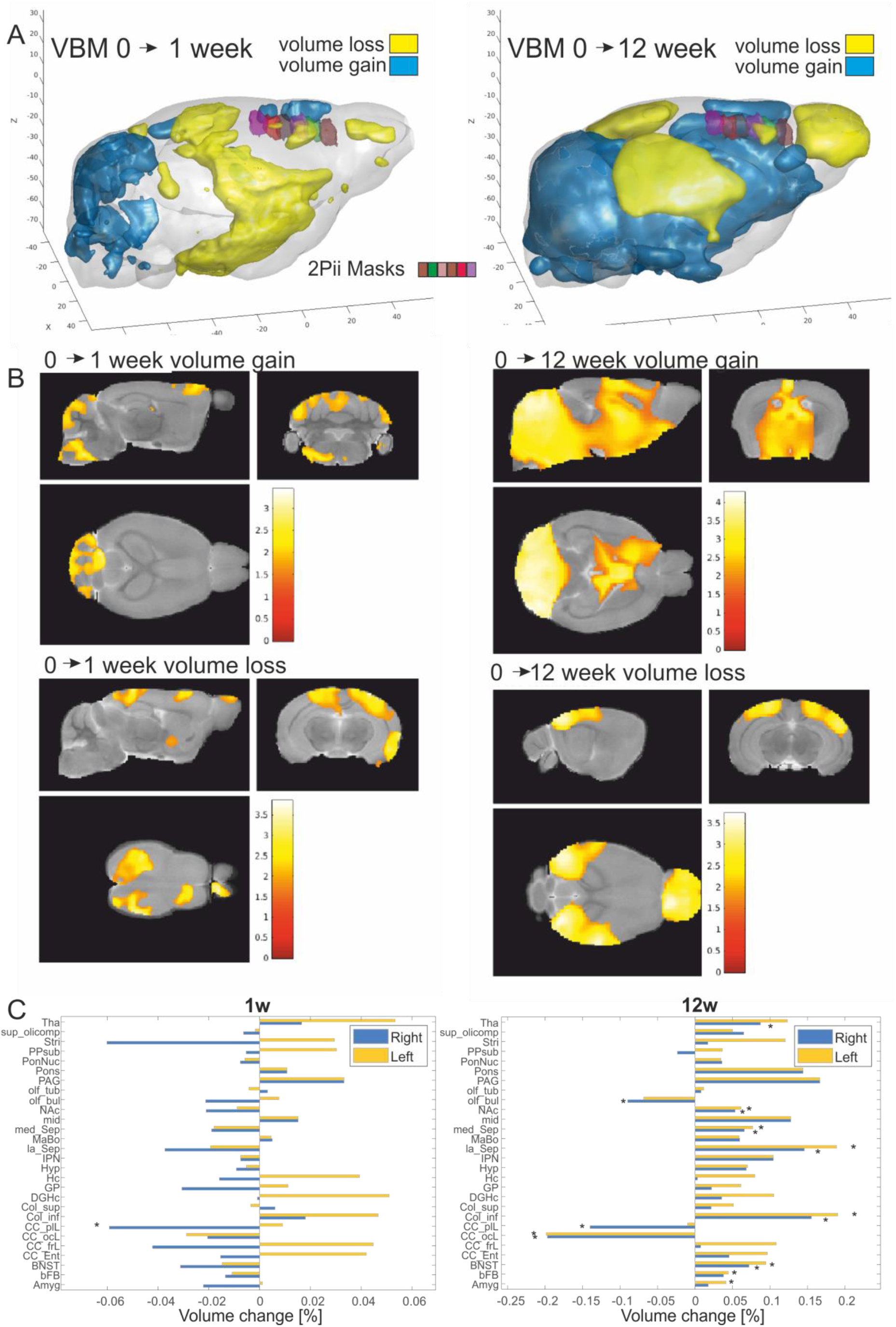
Longitudinal assessment of brain volume changes with VBM. **A.** Three-dimensional representation where positive (blue) and negative (yellow) volume changes were observed at 1 week and 12 weeks after baseline (p<0.05 uncorrected for display purposes). Imaging volumes addressed with 2Pii are labelled with distinct colours (see legend in left panel). **B.** Statistical maps depicting areas with significant volume growth and loss were detected (p< 0.05 uncorrected for display purposes, main clusters significant at p<0.05 cluster correction). **C.** Volume changes of 28 anatomical regions over time assessed by longitudinal VBM between the baseline measurement vs. one and twelve weeks. Regions with significant changes (p < 0.05) are marked (*). Abbreviations: Amygdala (Amyg), basal forebrain (bFB), bed nucleus of stria terminalis (BNST), entorhinal cortex (CC:Ent), frontal lobe (CC_frL), occipital lobe 130 164 (CC_ocL), parieto-temporallobe (CC_ptL), colliculus: inferior (Col_inf), colliculus superior (Col_sup), dentate gyrus of hippocampus (DGHC), globus pallidus (GP), hippocampus (HC), hypothalamus (Hyp), interpeduncular nucleus (IPN), lateral septum (la_Sep), mammillary bodies (MaBo), medial septum (med_Sep), midbrain (mid), nucleus accumbens (NAc), olfactory bulbs (olf_bul), olfactory tubercle (olf_tub), periaqueductal grey (PAG), pons (Pons), pontine nucleus (PonNuc), pre-para subiculum (PPSub), striatum (Stri), superior olivary complex (sup_olicomp) and thalamus (Tha).

### Imaging cell nuclei to derive structural parameters for superficial cortical volumes

To identify cell nuclei *in vivo*, we used mice genetically modified to ubiquitously express an EGFP-tagged Histone-H2B protein (‘Histone-GFP’, see methods section). The extent of labelling was tested by staining fixed brain sections with DAPI, an established reagent to stain DNA. Confocal imaging of Histone-GFP and DAPI revealed co-labelling of close to 100%, suggesting that in Histone-GFP mice almost all nuclei were labelled (Fig. 4A-C).

**Figure 4.**
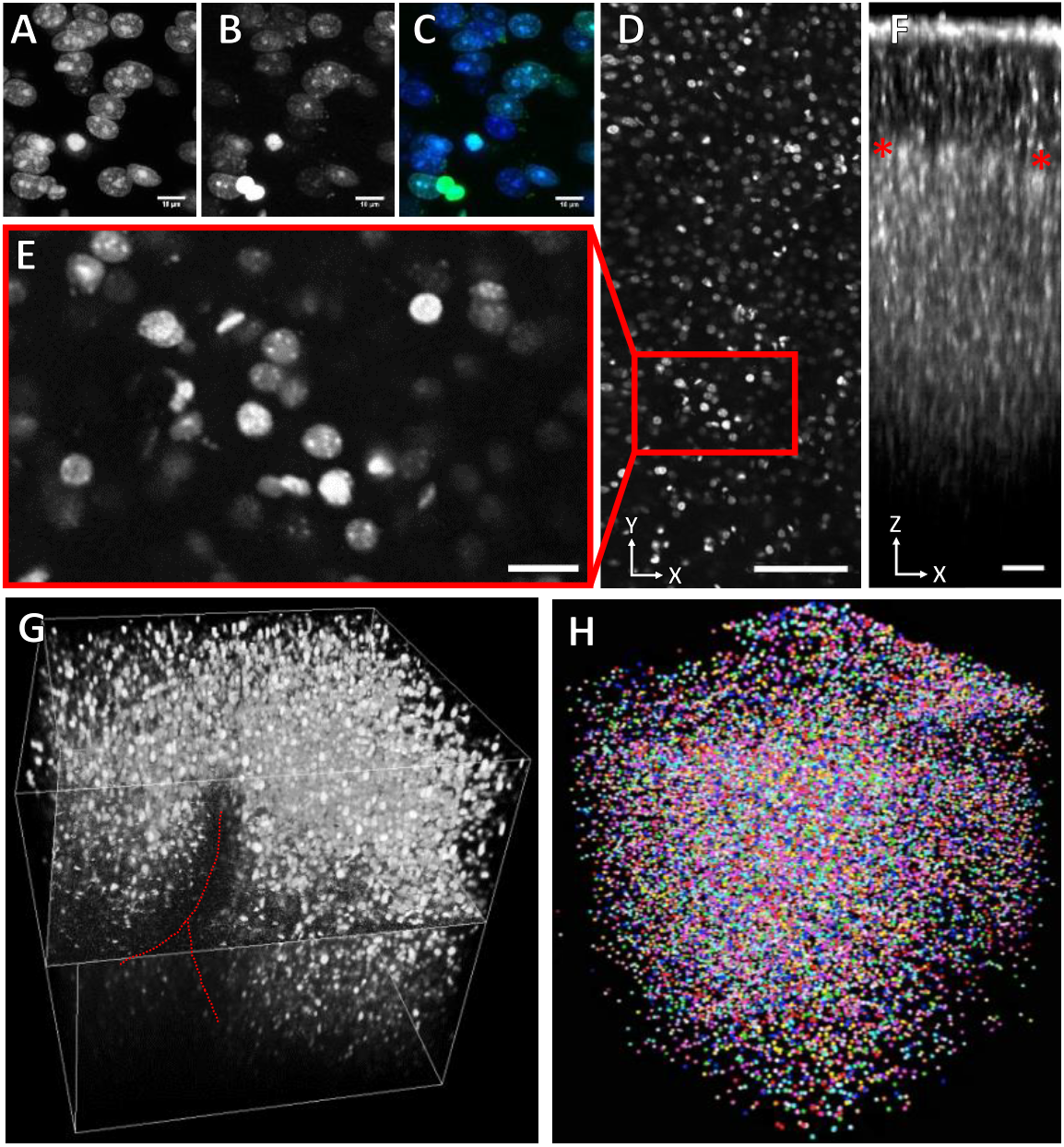
Histone-GFP labelled cell nuclei. **A-C.** Confocal Microscopy of fixed brain slices of Histone-GFP mice shows that all DAPI-stained nuclei **(A)** are also labelled with Histone-GFP **(B)** as seen in the overlap (**C**, green = Histone-GFP, blue= DAPI). **D.** Representative example image consisting of a maximum intensity projection of 10 consecutive image frames covering 20 µm of tissue recorded at a depth of 200 µm in Histone-GFP using 2Pii. Scale bar = 100 µm. **E.** Close up of D. Scale bar = 20 µm. **F.** Average intensity projection of a stack cropped in zx. The surface of the brain is located at the top of the image. Red stars indicate delineation of layer borders. Scale bar = 50µm. **G.** A representative 700 µm x 700 µm x 700 µm stack recorded by in vivo 2Pii in a Histone-GFP mouse. One orthogonal slice is displayed in the middle of the stack. The red stippled line denotes the position of a blood vessel. **H.** Segmented nuclei of a stack volume randomly coloured. Each sphere corresponds to the centroid position of a nucleus.

A representative image recorded with 2Pii through the cranial window at a depth of 200 µm shows densely packed nuclei (Fig. 4D). The magnified view highlights that nuclei were often sub-structured and of different size and shape (Fig. 4E). A typical image stack reveals layers differing in the density of nuclei, consistent with the layered structure of the neocortex (Fig. 4F). The most superficial densely packed nuclei are likely to be part of the pial membrane and astrocytes contributing to the glial limiting membrane. The border between the molecular layer and layer 2/3 could be readily identified (asterisks in Fig. 4F). A 3D projection of a complete image stack covering 700 µm x 700 µm x 700 µm shows the full extent of the volume acquired with one stack in 2Pii (Fig. 4G). For each microscopy session, six stacks were acquired, covering part of the medial right hemisphere (see Figs 2C, 3A). The example shown in Fig. 4G illustrates that a large fraction of nuclei residing within the imaging volume can be detected. However, limitations in acquiring images with high signal to noise ratio exist in deep neocortical areas (typically below 500 µm) and below larger blood vessels (dashed red line in Fig. 4G).

In summary, these results show that 2Pii of nuclei in neocortical areas corresponding to defined MRI volumes is feasible and delivers image data qualitatively suitable for several types of analyses as further outlined below. For example, these image data were processed by an automated pipeline (see Methods) and resulted in a 3D matrix of centroid positions each reflecting the location of a nucleus (Fig. 4H).

### Physical volume can be inferred from convex hull volumes spanned by identified marker nuclei

To identify changes of physical cortical volume from the positions of cell nuclei within a 2Pii stack, we developed a method considering several important confounding factors. On the timescale of the imaging intervals, the position of a cell could not be assumed to be stable, because cell death, proliferation, and migration may change the composition and geometrical arrangement of nuclei without affecting the local tissue volume. To circumvent these problems, we manually searched for nuclei that could be unequivocally identified at all three timepoints investigated. When inspecting the 2Pii stacks, we found that many nuclei did in fact stay in very recognizable, stable local patterns with other neighbouring nuclei over many months (Figure 5A). As it seems unlikely that this whole pattern of cells actively migrates together, we assumed that a shift in the relative stack position of a nucleus that stays in such a stable pattern can only come about by a shift in the volume around that pattern. A concerted migration of a subset of nuclei into different directions seems highly unlikely. Following this rationale, we manually sampled the stack positions of the same subset of nuclei (typically >30 to obtain reliable results) at all experimental time points (Fig. 5B). From this subset of cortical nuclei, distributed across the volume of interest, the surface formed by connecting nuclei using a 3D Delaunay triangulation was defined and subsequently the volume of the resulting convex hull calculated (see methods section and Fig. 5C). The convex hull could then be readily compared over all three timepoints, indicating an overall shrinkage or expansion. We sampled convex hulls in 1-2 of the 6 imaging positions in every animal along the rostrooccipital axis of the medial right hemisphere (see Fig. 3A, 2C). The volume of the convex hulls was in the range of 0.04 to 0.13 mm3 and was defined by the arbitrary choice of identified nuclei rather than reflecting the total reference volume.

**Figure 5.**
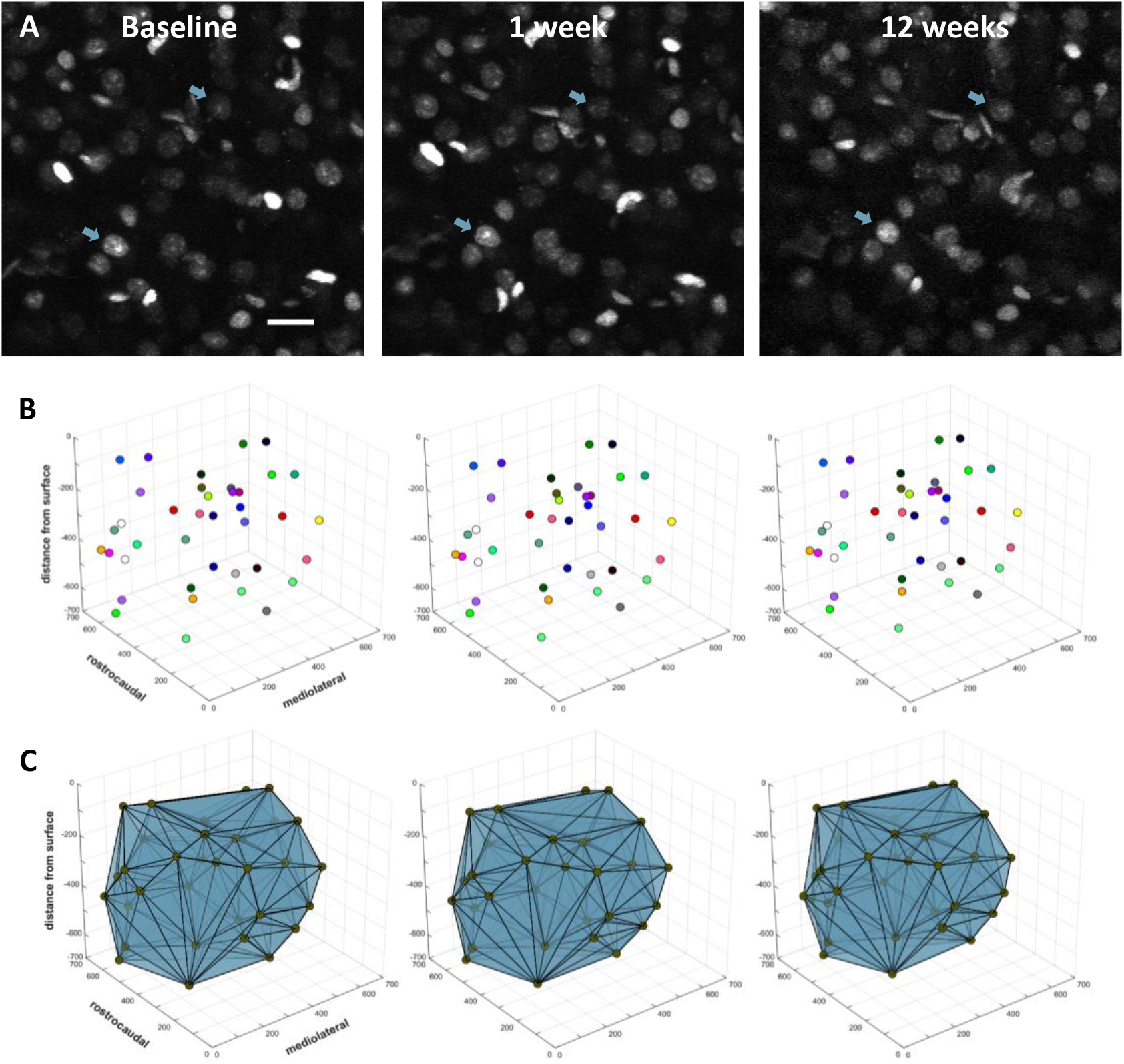
Determination of physical tissue volume from nearest-neighbour distances of identified nuclei. **A.** Unchanging patterns of nuclei were re-identified over all timepoints and nucleus centre coordinates were sampled as fiducials throughout all stack dimensions. Two examples of re-identified nuclei are indicated by blue arrows. Maximum intensity Z-projection of 10 optical sections covering 20 µm. Scale bar = 20 µm. **B.** Fiducial coordinates plotted inside single stack volume. Dot colours specify fiducials corresponding over time. All numbers in µm. **C.** Three-dimensional Delaunay triangulation from fiducials create matching tetrahedra whose subtle volume changes indicate shrinkage or expansion of the tissue within.

### GMV changes do not correlate with physical volume changes

The convex hull volumes determined for 9 mice at each timepoint revealed a statistically significant decrease from week 0 to 12 and from week 1 to 12 (Fig. 6A). At week 12, the volume decreased to approximately 94% of baseline (Fig. 6B), suggesting a shrinkage of the tissue by 6%. When analysing the corresponding GMV changes within MRI masks corresponding to stacks with a determined convex hull, we found a decreased GMV relative to baseline at week one, but not at week 12, with no significant difference between week one and 12 (Fig. 6C). Finally, the changes in GMV were not correlated with the changes in convex hull volume (Fig. 6D), indicating that the physical volume of brain tissue does not contribute to GMV determined with VBM in a significant way.

**Figure 6.**
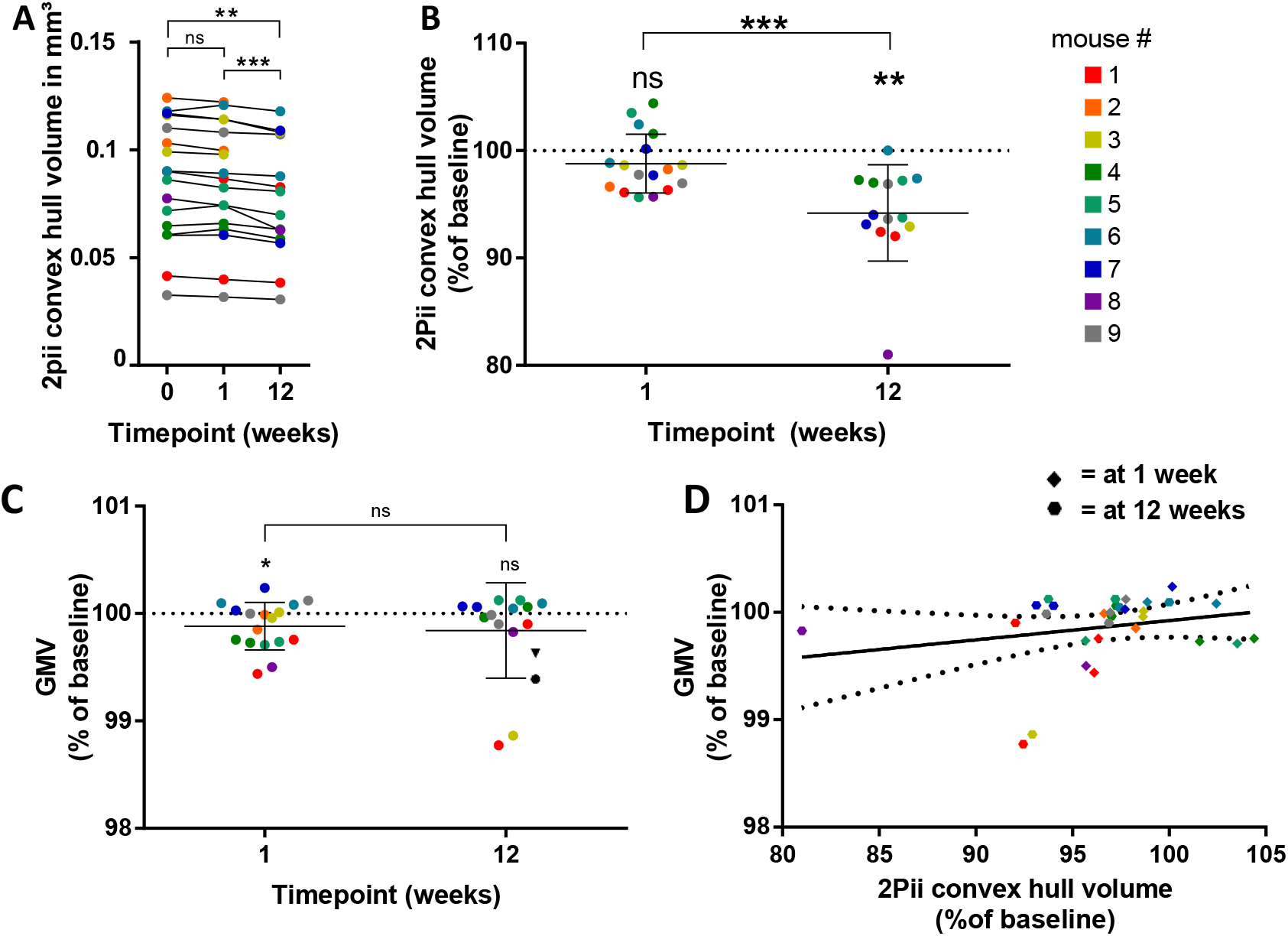
Correlation of physical volume with GMV. **A.** The volumes of convex hulls in 2Pii stacks significantly shrink over time. Display of all measured absolute volumes. Repeated measure one-way ANOVA with post-hoc Tukey test for multiple comparisons was done with complete datasets of n = 14 positions in 8 mice. Adjusted p values: 0 vs 1 week: p=0.4243; 0 vs 12 weeks: p = 0.0022; 1 vs 12 weeks: p= 0.0006. **B.** Relative change in 2Pii convex hull volumes compared to baseline at 1 week and 12 weeks. Same data and statistical test as in (A). At 1 week: mean ± SD = 98.79 ± 2.74 %; at 12 weeks: mean ± SD = 94.20 ± 4.50 %. Error bars indicate mean and standard deviation. **C.** Changes of GMV are plotted for positions that correspond to the 2Pii stacks in which the convex hull volume was defined. Percentages of signal are given as compared to baseline. Data were tested with one-sample t-tests against baseline and paired t-test between 1 and 12 weeks. At 1 week: n = 17 positions of 9 animals, mean/SD = 99.88%/0.22; 1 week compared to baseline: p = 0.0440. At 12 weeks: n= 14 positions of 8 animals, mean/SD = 99.84%/0.44; 12 weeks compared to baseline: p = 0.2061. 1 week vs 12 weeks: p = 0.8260. Error bars indicate mean and standard deviation. **D.** Microscopic volume changes do not explain changes in VBM. Linear regression analysis with R^2^=0,05223, p= 0.2163, n = 31 (16 positions of 9 animals at 1 week, 14 positions of 9 animals at 12 weeks). Solid line shows regression equation Y = 2.914*X – 194.3, dashed lines indicate 95% confidence intervals. Colour code denotes individual mice (see Fig. 6B).

### Correlations of GMV with the number of cells

We next tested if changes in the number of nuclei within a 2Pii stack correlated with alterations in the respective GMV. The number of nuclei and all subsequent readouts were derived from our automated image analysis for 3D nucleus detection described above (see Methods, Fig. 4H), which allowed us to investigate more datasets than we did for manual determination of physical tissue volume. Consequently, 28 matching MRI masks were used for GMV calculation and correlation. Based on this analysis, GMV was significantly decreased at week 12, both in comparison to baseline and week one (Fig. 7A).

**Figure 7.**
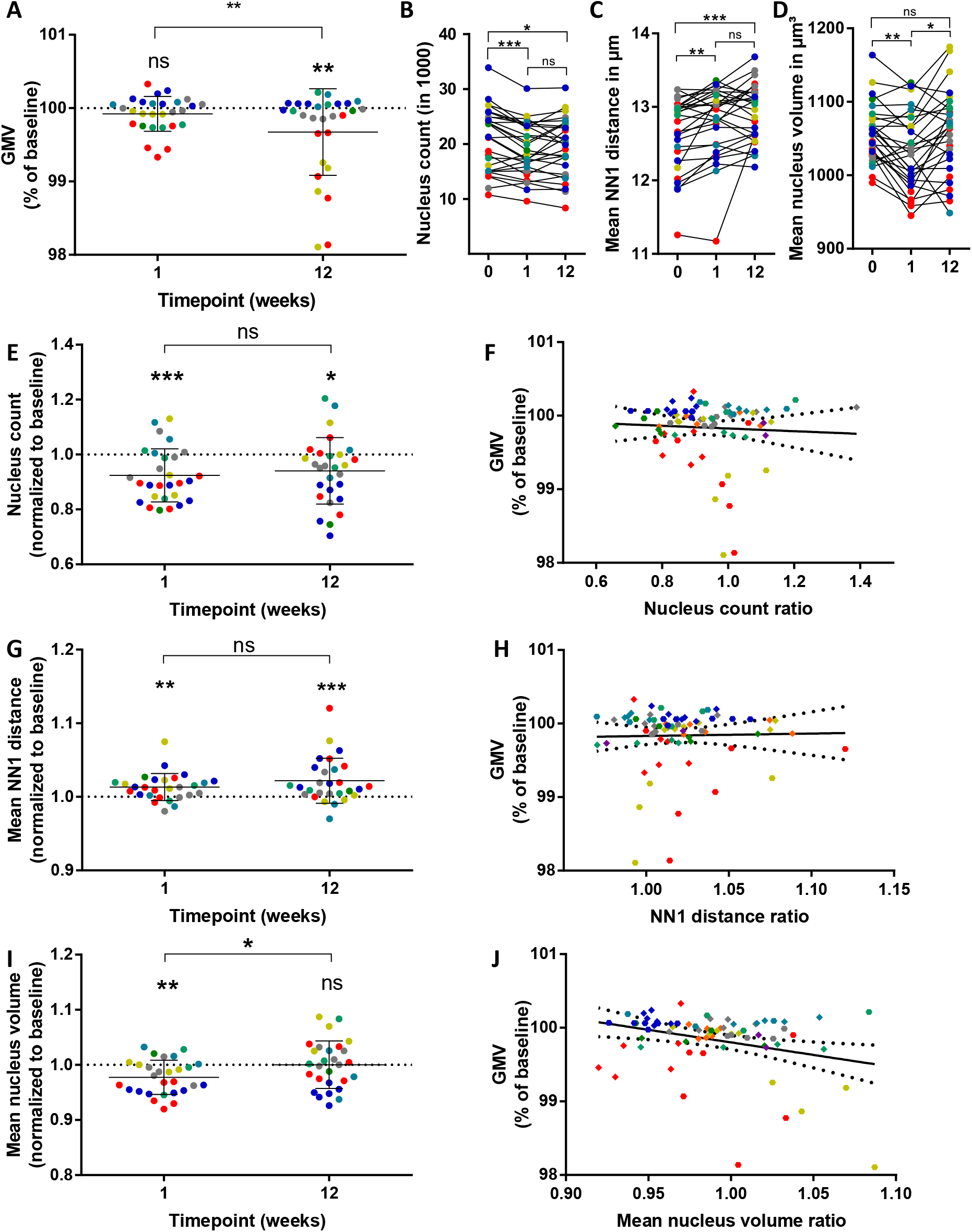
Correlations of GMV with cell number, nearest neighbour distance and mean nucleus volume. **A.** GMV changes in volumes matching to 2Pii stacks that were studied by automated analyses. Data were tested with one-sample t-tests against baseline and paired t-test between 1 and 12 weeks, n = 28 positions of 7 animals. At 1 week to baseline: p=0.0942; at 12 weeks to baseline: p = 0.0069; at 1 week to 12 weeks: p = 0.0016. Mean ± SD in % at 1 week = 99.92±0.24; at 12 weeks = 99.67±0.59. Error bars indicate Mean ± SD. **B.** Number of cells as revealed by nucleus count. Repeated measure one-way ANOVA with post-hoc Tukey test for multiple comparisons, n = 28 positions in 7 animals. Mean ± SD (in 1000) at baseline = 20.78±5.73; at 1 week = 19.00±4.87; at 12 weeks = 19.40±5.25. Adjusted p-values: 0 vs 1 week: p = 0.0002; 0 vs 12 weeks: p = 0.0239, 1 vs 12 weeks: p = 0.6286 **C.** Mean nearest neighbour distance. Friedman test with post-hoc Dunn’s test for multiple comparisons was done with n = 28 positions in 7 animals. Median/interquartile range in µm at baseline = 12.85/12.30-13.01; at 1 week = 12.97/12.48-13.19; at 12 weeks = 13.00/12.62-13.19. Adjusted p-values: 0 vs 1 week: p = 0.0040; 0 vs 12 weeks: p = 0.0009, 1 vs 12 weeks: p > 0.9999. **D.** Mean nuclear volume. Repeated measure one-way ANOVA with post-hoc Tukey test for multiple comparisons was performed with complete datasets of n = 28 positions in 7 animals. Mean ± SD in µm^3^ at baseline = 1057±39; at 1 week = 1033±52; at 12 weeks = 1057±58. Adjusted p-values: 0 vs 1 week: p = 0.0040; 0 vs 12 weeks: p = 0.0009, 1 vs 12 weeks: p > 0.9999. **E.** Relative change in the number of cells. Same data and statistical test as in (B). Line and error bars indicate Mean ± SD; at 1 week: 0.924±0.097; at 12 weeks 0.941±0.121. **F.** Correlation of GMV and number of cells. Linear regression with n = 74 from 9 animals and 2 timepoints, p=0.6349. **G.** Relative change in nearest neighbour distances. Same data and statistical test as in C. Line and error bars indicate Mean ± SD; at 1 week: 1.013±0.018; at 12 weeks 1.022±0.031. **H.** Correlation of GMV with NN1 distance. Linear regression with n = 74 of 9 animals and 2 timepoints, p = 0.8516. **I.** Relative change in mean nucleus volume. Same data and statistical test as in D. Line and error bars indicate Mean ± SD; at 1 week: 0.978±0.031; at 12 weeks 1.000±0.043. **J.** Correlation of GMV and mean nucleus volume. Linear regression with n = 74 of 9 animals and 2 timepoints, p=0.0076, R^2^=0.0947, Equation: Y = −3.410*X+103.2. Dashed lines in plots F, H and J represent the 95% confidence band for the regression.

The 3D matrices of nucleus positions (Fig. 4H) revealed on average 20000 nuclei per volume (Fig. 7B), with a pronounced variability between areas and mice. This can be explained by the different extent and size of blood vessels contained in the imaging volumes (see Fig. 4G). The number of nuclei decreased significantly from baseline to week 1 and week 12, however, week 1 and 12 were not distinguishable. The number of nuclei decreased by approximately 8% relative to baseline, suggesting a substantial amount of cell loss after performing the baseline measurement. Surprisingly, these changes in cell number were not correlated with GMV changes (Fig. 7F), indicating that even pronounced changes in cell number do not contribute to GMV.

### Correlations of GMV with nearest neighbour distances

In addition to the physical volume and cell number we determined the nearest neighbour distance (NN1), reflecting the distance between the detection of each nucleus to the detection of the nucleus nearest to it in Euclidean space. The mean NN1 of all NN1 in a stack were then calculated. Changes in NN1 may be induced by different mechanisms acting in parallel (Fig. 1D): (1) Uniform expansion or shrinkage of local tissue volume, with tissue expansion generating larger NN1; (2) Gain or loss of cells in the immediate space surrounding a nucleus. For example, loss of the nearest neighbour would increase NN1. (3) A different distribution of nuclei, for example local clustering would decrease NN1. The mean NN1 at baseline was 12.65 ± 0.5 µm, with an increase to 12.81 ± 0.48 µm after one week and 12.91 ± 0.39 within 12 weeks (Fig. 7C). Hence, a small increase of approximately 2 % in NN1 could be detected (Fig. 7E). This increase is consistent with and could thus be explained by the decreased cell number reported above (Fig. 7B). Correlating NN1 to GMV did not reveal a significant relationship (Fig. 7H).

### Correlations of GMV with mean nucleus volume

Finally, we determined the mean volume of all nuclei. With this measure we intended to gain additional information about the respective cells. A change of observed mean nucleus volumes could mean a (1) change in relative amount of cell types within the investigated cortical volumes as e.g. glial cells tend to have smaller nuclei than neuronal cells, or (2) a change in nucleus size is a direct indicator for different transcriptional activity states (Fair, Hyttel et al. 1995). We found a significant decrease of nuclear volume from baseline to week one by approximately 2%, followed by a significant increase to week 12 (Fig. 7D, 7I). The average volume at baseline and week 12 was indistinguishable. These changes in nuclear volume were inversely correlated with the GMV (Fig. 7. J). Hence, a decrease in GMV is accompanied by an increase in mean nuclear volume.

### 2Pii parameters correlate with GMV in certain cortical depths

The findings above were results generalized for the whole 2Pii stack volume. Since the cerebral cortex is organized in specified layers, it seems possible that mechanisms influencing GMV might be different in different cortical depths. We addressed this question by investigating 2Pii parameters derived in confined stack sizes of 700 µm x 700 µm x 175 µm (XYZ) each that correlate with the GMV change in the MRI mask. A higher nucleus count within the surface and layer I region of the cortex (0-175um) was associated with a higher GMV (Fig. 8A). Similarly, the mean NN1 correlates significantly in the same layer, again hinting at the nucleus density being the main factor influencing NN1 (Fig. 8A). The layer which best explained the variation in GMV by nucleus volume was in a deeper layer, between 350 and 525 µm (Fig. 8C).

**Figure 8.**
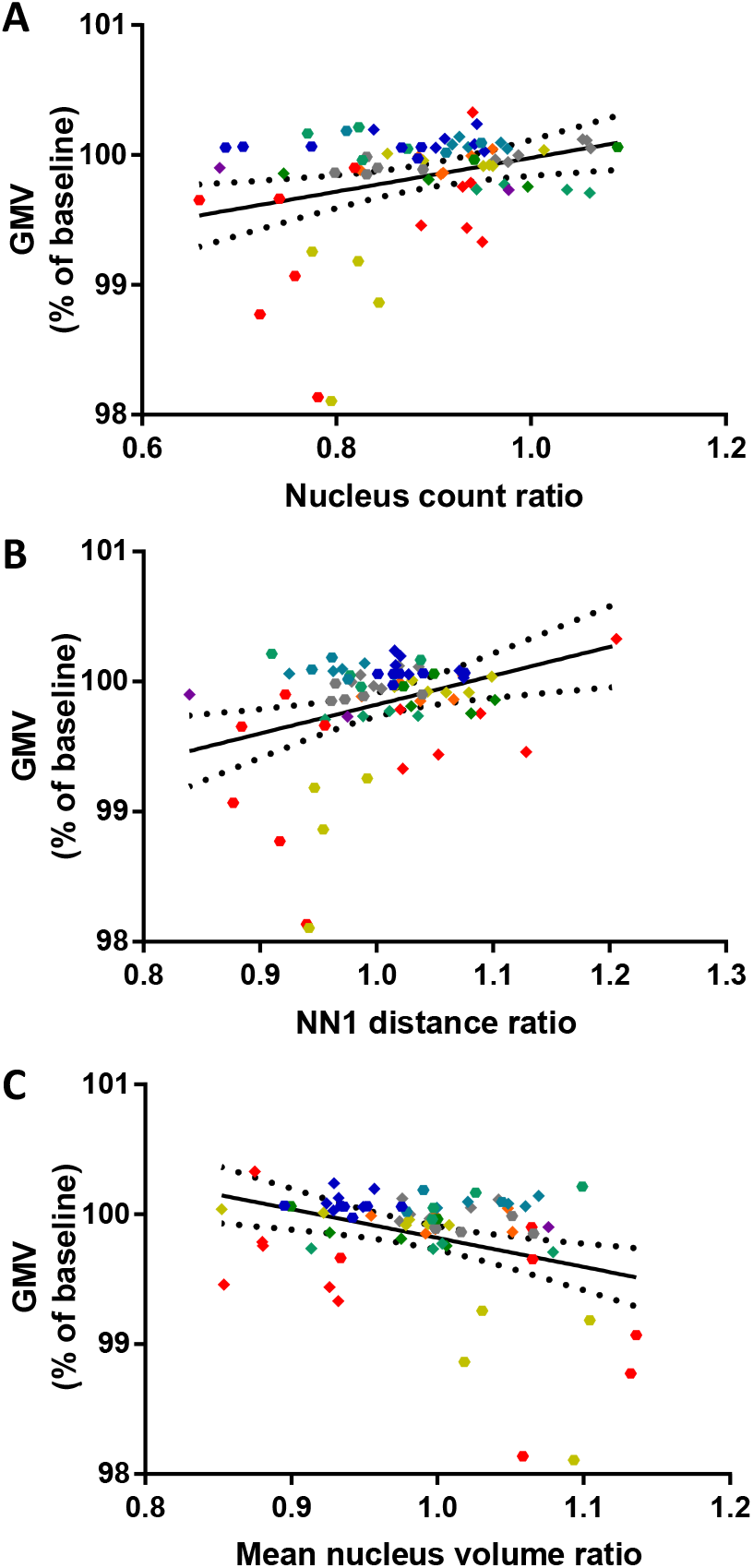
Correlations within sub-layers of the 2Pii data. **A.** Correlation of GMV changes and nucleus count limited to the z-range from 0 to 175 µm from the cortical surface. Linear regression with n=74 of 9 mice and two time points, p = 0.0081, R^2^ = 0.0932, equation: Y = 1.301 * X + 98.68. **B.** Correlation of GMV changes and NN1 distance limited to the z-range from 350 to 525 µm from the cortical surface. Linear regression with n=74 of 9 mice and two time points, p = 0.0053, R^2^ = 0.1031, equation: Y = 2.221 * X + 97.60. **C.** Correlation of GMV changes and mean nucleus volume limited to the z-range from 350 to 525 µm from the cortical surface. Linear regression with n=74 of 9 mice and two time points, p = 0.0026, R^2^ = 0.1195, equation: Y = −2.226 * X + 102.0.

## Discussion

This study combined small animal MRI with alternating 2Pii to systematically correlate VBM measures with cellular composition. The longitudinal design delivered intra-individual comparisons at different time points in a paradigm based on age-dependent changes in GMV. We found that a decrease in physical volume did not correlate with GMV, while the NN1 and nuclear volume did show correlations. Restricting the correlations to sublayers of cellular imaging data, these correlations became stronger for NN1 and nuclear volume and significant for GMV and cell count. Nevertheless, these correlations explain only a small fraction of changes observed in GMV changes. Thus, our study revealed that GMV changes are not dominated by changes in physical volume and that local cell distribution, cell number and nuclear volume contribute.

### Technical aspects

Our approach compares GMV and cellular organization in the native state using a longitudinal design. While this represents a major advantage, several technical issues may limit the conclusions of our study: (1) Effects of chronic cranial window on VBM. The implanted windows, while necessary for high fidelity 2Pii, may affect the MRI signal and thus impact the VBM procedure. We found a change in the physical boundaries of the brain in the frontal area that we could improve by using curved windows, closely following the anatomical shape of the brain. Furthermore, the window surgery may exert local effects on the underlying tissue. In our T2-weighted anatomical MR images we found hyperintense areas close to the edges of the window in some mice that we attribute to CSF accumulation, local oedema or scar tissue formation along the rim of the window. On the other hand, these areas were outside of the brain and could be eliminated by the automatic brain extraction algorithm. In summary, we think that these effects do not have a major impact on our VBM analysis, supported by the finding that the volume changes described here are within the range found in previous studies (Bilbao, Falfan-Melgoza et al. 2018). (2) Effects of anaesthesia. In both imaging modalities, the same anaesthesia was applied, hence providing comparable conditions. (3) Cross-modal registration. Errors in registration may affect the outcome of correlations. However, the extent of unprecise registrations would not impact our correlations much. (4) Reliability of cell counting. The counting algorithm itself can be considered highly reliable. However, the presence of blood vessels in the imaging volume can lead to differences in the number of nuclei detected within the six different imaging volumes. This includes volume-displacement of the blood vessel itself and shadowing effects on nuclei positioned underneath the vessel (typically at deeper z positions). Thus, we cannot claim that all nuclei of the imaged volume can be determined, yet a reproducible number at different time points and a number large enough to warrant the analyses performed here can be identified.

Taken together, we conclude that the approach introduced here represents a valid and advantageous strategy to identify mechanisms contributing to GMV changes.

### Age-dependent changes in VBM

The grey matter changes detected by VBM follow a pattern previously described (Bilbao, Falfan-Melgoza et al. 2018), with age-related volume increasing in the cerebellum and subcortical areas (midbrain, hindbrain, thalamus, hypothalamus and pallidum/lateral septum), and reduced growth or loss of volume being present in the olfactory bulb and distinct cortical areas (visual cortex, somatosensory area and motor cortex as well as parts of the anterior cingulate). However, we found a stronger overall increase in volume in the previous study which can be attributed to the later starting age of the current study. The described changes are consistent with volume changes found in human aging (Good, Johnsrude et al. 2001). In conclusion, the paradigm used here recapitulates previous age-related changes, further underlining the validity of the MRI imaging approach in mice carrying cranial windows.

### Correlation of GMV with cellular metrics

Our study addressed four structural parameters that may each contribute to a certain extent to GMV changes:

(1) Physical volume. The actual change of physical volume within the tissue remains to be validated for the vast majority of cases where VBM has been applied. Keifer et al. (2015) have approached this question studying cortical thickness ex vivo in fixed brain slices. They did not find significantly thicker cortices in regions where ex vivo VBM had revealed GMV increases. In our study we introduced a 3-dimensional readout of physical volume changes by taking distances between reidentified cell nuclei as measurement for tissue volume. Although in VBM as well as 2Pii shrinkages were observed over time, both these volume measures did not correlate, consistent with the described findings by Keifer et al. (2015). It seems possible that the GMV readout is influenced less by real volume changes than by other factors that may account for GMV alterations. The absence of a direct correlation between the physical cortical volume calculated by the position of cell nuclei within a 2Pii stack and the VBM GMV changes might also be due to the difference in scale by the underlying algorithms that are employed during the longitudinal registration in VBM. Since the changes in the MRI volume are calculated by a comparably smooth function over the whole cortical thickness, the major changes might appear further away from the brain surface. Furthermore, changes in physical volume may come about via alterations of the extracellular space or changes in the size of the neurons and their compartments such as the dendritic and axonal trees. Previous reports suggested that neuronal structural plasticity may contribute to changes in GMV (Biedermann, Fuss et al. 2012, Suzuki, Sumiyoshi et al. 2013, Keifer, Hurt et al. 2015) (Biedermann, Keifer, Suzuki). For example, Keifer et al. (2015) proposed that a higher spine density explains 20 % of the increased ‘VBM signal’ in the auditory cortex of mice after auditory fear conditioning compared to a control group (Keifer, Hurt et al. 2015). These studies used a cross-sectional design and determined microscopic correlates in fixed brain slices *ex vivo*. It seems likely that the procedure of brain fixation with paraformaldehyde (PFA) may affect the volume and shape of the brain (Wehrl, Bezrukov et al. 2015, Hikishima, Komaki et al. 2017) with some areas being more affected than others. Furthermore, the cross-sectional design may lack sensitivity and statistical power.

(2) Cell number. Although we found a decrease in cell number after one and 12 weeks, this decrease did not correlate with changes in GMV when comparing entire imaging volumes covering cortical layers 1 to 5. The layered organization of the cortex prompted us to limit correlations to smaller sub-volumes corresponding to different cortical layers. With this approach, a significant correlation of GMV changes with the cell count could be detected in the most superficial layer (Fig. 8A). This finding suggests that changes in cell number within limited areas may contribute to GMV changes. Hence, mechanisms leading to cell death, neurogenesis, astrogliosis, and microglial migration need to be taken into consideration. Often, these processes are organized in circumscribed areas, rather than being distributed across entire neocortical layers.

(3) Nearest neighbour distance (NN1). When looking at correlations of the NN1 with GMV within the entire 2Pii volume, no significant relation could be detected. However, when limiting the analysis to deep cortical layers, changes in NN1 were significantly correlated with changes in GMV. Again, this may indicate that a more localized process can affect the GMV readout. We interpret the NN1 such that positional changes do not occur in terms of local clustering (see Fig. 1D), but rather indicate an effect evenly distributed across the volume analysed.

(4) Nucleus volume. The mean nucleus volume was the only parameter that inversely correlated with GMV when analysing entire matched imaging volumes. When focusing on deeper layers, this correlation improved. While the mechanistic basis and consequences of smaller or larger nuclei on GMV remains unclear, we speculate that transcriptional activity in general may affect the size of the nucleus. Nuclei with a large fraction of transcription inactive heterochromatin are typically smaller than nuclei with a large fraction of euchromatin (e.g neurons, being the most transcriptionally active cells, have pronounced euchromatic nuclei that are typically much larger than astrocytic or oligodendrocytic nuclei; heterochromatic DNA is more densely packed).

## Conclusions

Our study provided a systematic assessment of several structural parameters and their contribution to changes in GMV. We could show that GMV changes are not much related to changes in physical volume, suggesting that voxel-based volume shifts cannot be interpreted as shifts in physical volume. A large range of mechanisms remains that may contribute to these changes. Our work proposes that local cell count, spatial arrangement of somata and their nuclei as well as nuclear volume are factors that may contribute to some extent. Furthermore, the results suggest that spatially restricted processes may influence GMV changes derived from larger areas. We predict that the main contributions are provided by the structures residing between the nuclei, consisting of neuropil, glia and vasculature. The details remain to be figured out, the correlative *in vivo* approach introduced here, extended by additional readouts such as dendritic, glial and vascular compartments, as well as MRI markers like fractional anisotropy and changes in relaxation parameters, may support the quest to understanding the mechanisms underpinning GMV changes.

## Acknowledgements

We want to thankfully acknowledge the Michaela Kaiser, Claudia Kocksch and Felix Hoerner for their excellent technical support during preparation and execution of experiments. The data storage service SDS@hd supported by the Ministry of Science, Research and the Arts Baden-Württemberg (MWK) and the German Research Foundation (DFG) through the grant INST 35/1314-1 FUGG and the high performance cluster bwForCluster MLS&WISO supported by the MWK are gratefully acknowledged. We also acknowledge the support of the DFG within the SFB1158 grants B04 (WWF) and B08 (TK).

## Contributions

L.A. performed 2Pii, animal surgery, analysed data and wrote the manuscript. C.B. developed the pipeline for automatic segmentation of the nuclei. W.W.-F. and C.F.-M. Performed MRI measurements and analysed MRI-Data. J.K., L.A., C.F.-M, W.W.-F. and T.K. wrote the manuscript. T.K. conceptualized the project, J.K. conceptualized and implemented the project, analysed data and wrote the manuscript. All authors reviewed and approved the manuscript.

---

**Supplementary Figure 1:**
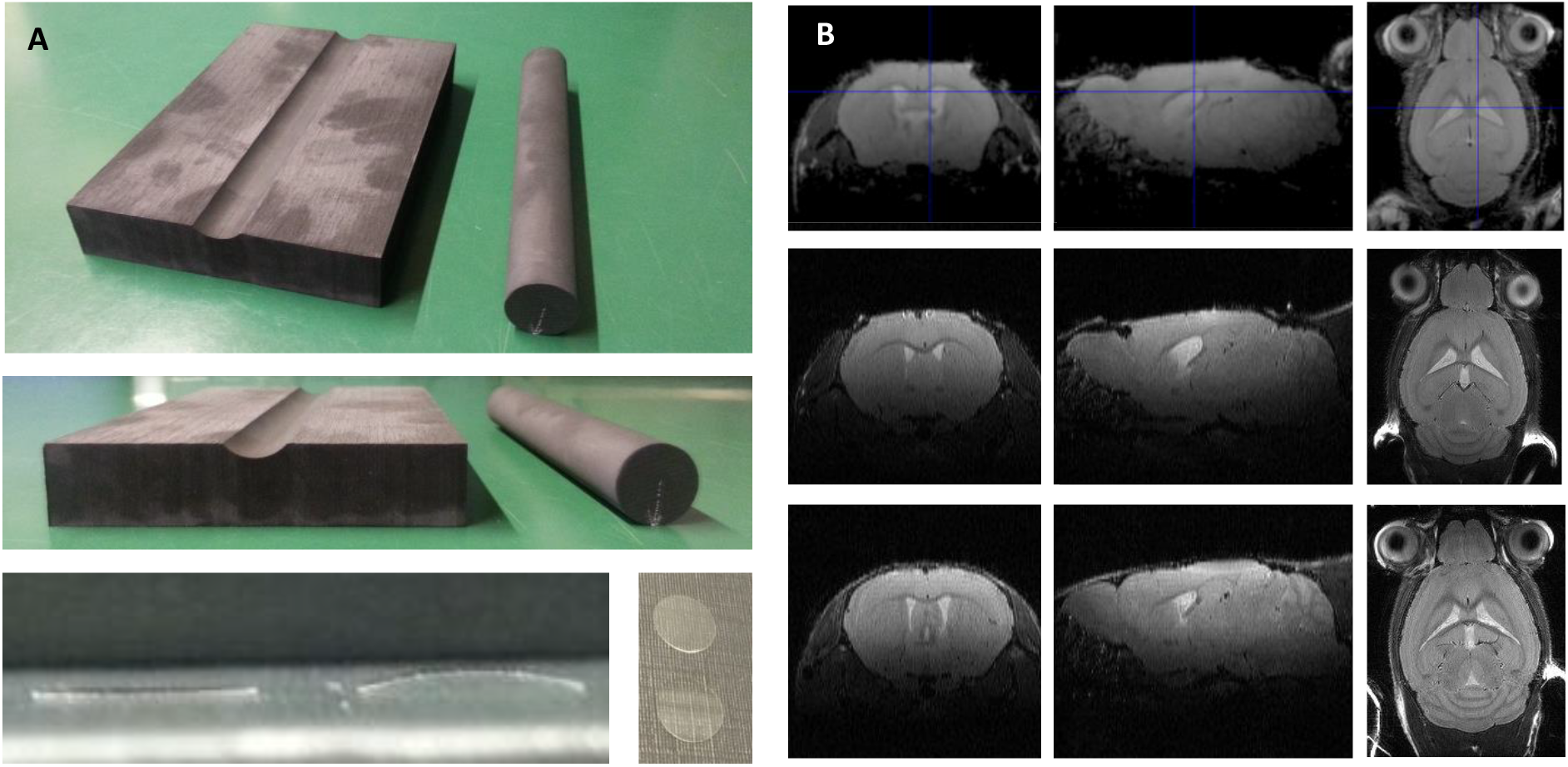
Glass coverslips for cranial window were curved to match the natural brain curvature. A. Bending the coverslips. A hollow was milled into a graphite block to match a 20 mm diameter graphite cylinder. Flat coverslips were lined up in the hollow, cylinder was placed on top and construct was heated in a laboratory oven. B. Comparing MRI scans of mice with a flat coverslip (top row), a bent coverslip (middle row) and a mouse without cranial window (bottom row). Less budging of the brain at the craniectomy and a more natural curvature is achieved by the bent coverslip.

**Supplementary Figure 2:**
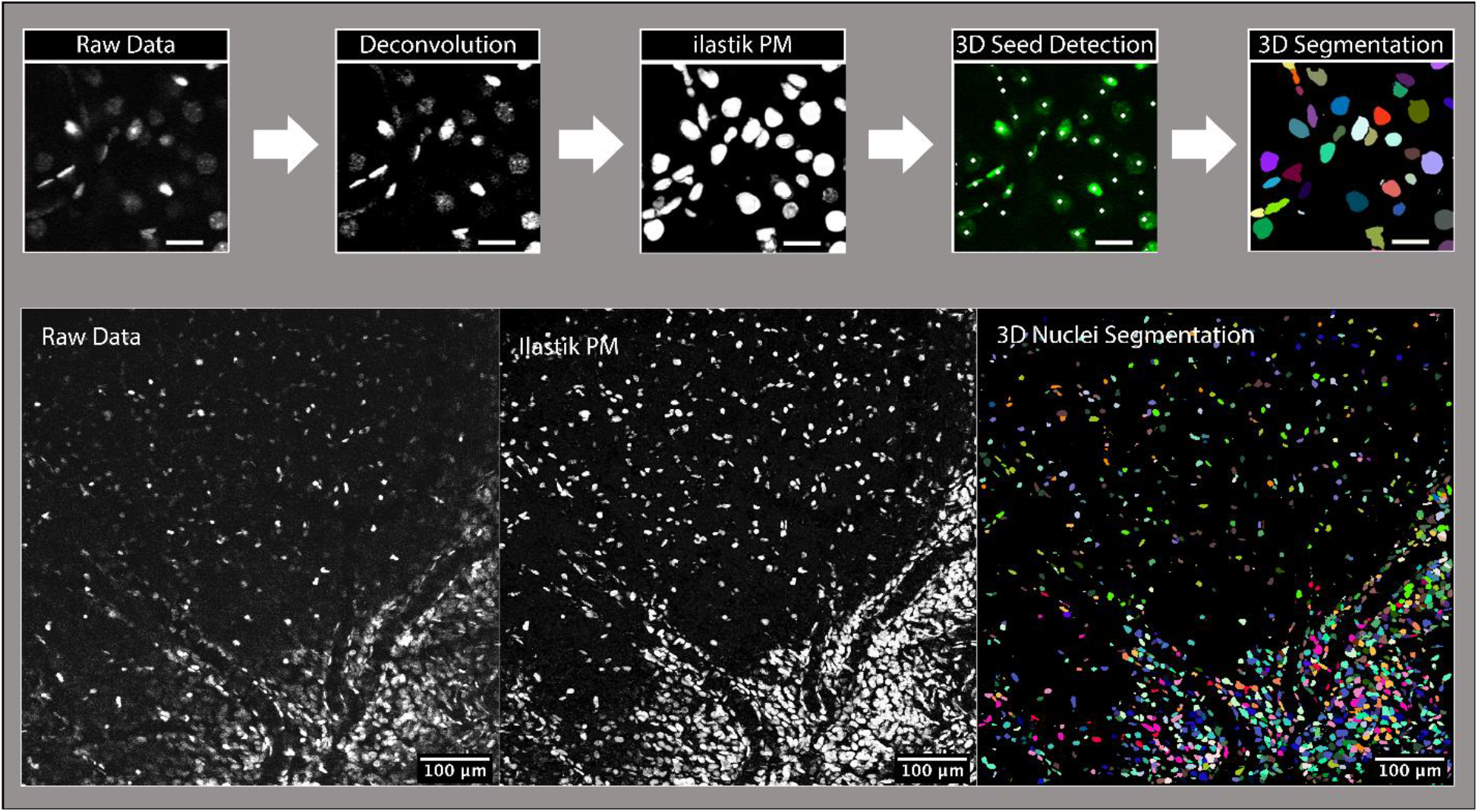
Analysis workflow for nucleus segmentation. See methods for a detailed description.

**Supplementary Movie 1: Visualization of the workflow for nucleus segmentation**

The 3D rendering was created using arivis Vision4D software version 3.0.1 (arivis AG (Munich, Germany). Raw data is displayed in green and probability map in grey. The centroids were imported as xyz coordinates from the output of the automated segmentation and visualized using random colours.

